# Impacts of competition and phenotypic plasticity on the viability of adaptive therapy

**DOI:** 10.1101/2025.03.20.644475

**Authors:** B. Vibishan, Paras Jain, Vedant Sharma, Kishore Hari, Claus Kadelka, Jason T. George, Mohit Kumar Jolly

## Abstract

Cancer is heterogeneous and variability in drug sensitivity is widely documented across cancer types. Adaptive therapy is an emerging modality of cancer treatment that leverages this drug resistance heterogeneity to improve therapeutic outcomes. Current standard treatments typically eliminate a large fraction of drug-sensitive cells, leading to drug-resistant relapse due to competitive release. Adaptive therapy aims to retain some drug-sensitive cells, thereby limiting resistant cell growth by ecological competition. While early clinical trials of such a strategy have shown promise, optimisation of adaptive therapy is a subject of active study. Current methods largely assume cell phenotypes to remain constant, even though cell-state transitions could permit drug-sensitive and -resistant phenotypes to interchange and thus escape therapy. We address this gap using a deterministic model of population growth, in which sensitive and resistant cells grow under competition and undergo cell-state transitions. Based on the model’s steady-state behaviour and temporal dynamics, we identify distinct balances of competition and phenotypic transitions that are suitable for effective adaptive versus constant dose therapy. Our data indicate that under adaptive therapy, models with cell-state transitions show a higher frequency of fluctuations than those without, suggesting that the balance between ecological competition and phenotypic transitions could determine population-level dynamical properties. Our analyses also identify key limitations of applying phenomenological models in clinical practice for therapy design and implementation, particularly when cell-state transitions are involved. These findings provide an overall perspective on the relevance of phenotypic plasticity for emerging cancer treatment strategies using population dynamics as an investigation framework.

**Significance Statement:** Drug-sensitive and -resistant cancer cells can compete with each other within the same tumour, and adaptive therapy exploits this competition to control overall tumour growth. The fact that sensitive and resistant cell types can switch phenotypes complicates the implementation of adaptive therapy. Our ODE-based theoretical ecology framework shows that asymmetric competition favouring sensitive cells generally benefits therapy outcomes, while phenotypic plasticity is usually detrimental. Our model also provides cell population-level indicators that can help predict the underlying balance between competition and plasticity. Phenomenological models are limited by whether their parameters can be reliably identified given scarce data, and our study illuminates the scope of such models in understanding cancer population dynamics and the need for mechanistic modelling.

## 1 Introduction

Cancer treatment is constantly evolving, much like cancers themselves. One of the more recent subjects of this ongoing evolution concerns the problem of intra-tumour heterogeneity (Almendro et al., 2013, Gay et al., 2016). The fact that cancer cells vary widely in their genotype as well as phenotype poses a significant challenge as it implies that a subset of cancer cells in a growing tumour is invariably resistant to a given drug treatment. Treatment with this drug therefore effectively removes only the sensitive cancer cells, and could directly contribute to drug-resistant relapse of the tumour through competitive release of drug-resistant sub-populations. Adaptive therapy seeks to manage this problem using an ecological approach (Gatenby et al., 2009, Hansen et al., 2017, Hansen and Read, 2020), whereby lower doses of a drug alongside drug holidays are used to maintain a small, controlled population of drug-sensitive cells, which in turn limit the growth of drug-resistant cells due to competitive inhibition. The idea has spawned a considerable body of theoretical work (Enriquez-Navas et al., 2016, Cunningham et al., 2018, West et al., 2020, Aguadé-Gorgorió et al., 2021, Strobl et al., 2021, Gedye and Navani, 2022) as well as clinical trails to test its practical feasibility (Zhang et al., 2017, 2022). Despite some empirical evidence that adaptive therapy could in principle outperform standard-of-care maximum dose regimens (Enriquez-Navas et al., 2016, Zhang et al., 2022), its applicability is far from universal.

The design and implementation of adaptive therapy is most effective when some knowledge of the relative frequencies of drug-sensitive and -resistant cells is available, even though some proxies of the total population size have still proven to be sufficiently useful in this context Zhang et al. (2017). On the other hand, success with adaptive therapy strongly depends on the strength and direction of cell-cell interactions remaining roughly constant throughout the duration of the treatment period, which is unlikely to be satisfied in real-life treatment situations. Phenotypic plasticity is known to be widely prevalent across cancers and cancer cells are often able to switch between alternate phenotypes that are both stable under different contexts. Epithelial-to-mesenchymal transitions (EMT) and their inverse, mesenchymal-to-epithelial transitions, are arguably the best characterised instance of such cell-state switching in cancer (Kalluri and Weinberg, 2009, Bastid, 2012, Ye and Weinberg, 2015, Jolly et al., 2019), but such plasticity has been documented in other aspects of cell biology, including cell death and drug resistance (Sharma et al., 2010, Huang et al., 2013, Wang et al., 2024). In the context of adaptive therapy, plasticity is directly relevant to the ecological dynamics of the tumour population as a whole. In general, the strength of biotic inhibition exerted by a given sub-population is typically a direct function of its current abundance. This could be diluted by switching between the sensitive and resistant cell states, and thus cause non-linearity in how the overall strength of biotic inhibition scales with the abundance of each sub-population. It is also possible that sensitive cells could escape therapy by switching temporarily to a resistant state, and switch back after some period of time to recapitulate the same population distribution. Recently, a wealth of information from single-cell genomic and transcriptomic data analyses have significantly improved our mechanistic understanding of how phenotypic plasticity operates at the cellular level (Caiado et al., 2016, Almendro et al., 2013, Deshmukh et al., 2021). However, except for some early work (Shah et al., 2023), this understanding has yet to be extended to the population-level, where it would be directly relevant to the design and implementation of novel therapies. Moreover, while some recent work has addressed the dynamical effects of plasticity in a model of density-dependent population growth (Kim et al., 2021), a comprehensive integration of competition and plasticity within a single modelling framework is still lacking.

In this study, we address this gap by developing a simple model of a generalised two-cell type cancer system that includes (1) density-dependent competition, and (2) phenotypic plasticity in the form of cell-state transitions between drug-sensitive and -resistant cells. Using a modified logistic framework, we investigate the relationship between competition and transitions in the overall population dynamics of this system, first in the absence of therapy, and then under different dosing regimens of a cytostatic or a cytotoxic therapy. On the whole, our results illustrate a complex interplay between competition and phenotypic plasticity in determining the steady state behaviour of the system, particularly when fixed points cannot be identified directly. We expand this theoretical insight using numerical simulations, which identify a set of minimal conditions that represent a favourable balance between competition and transitions in terms of therapeutic outcomes. Our data also reveal differences in temporal patterns in the total population dynamics that could be used to infer the relative importance of competition vs. transitions in determining underlying behaviour. Finally, since our model is essentially phenomenological in nature, we use existing analytical methods to demonstrate a marked lack of structural identifiability of the parameters in such a framework. This paucity in turn limits the extent to which its parameters can be reliably estimated in a data-scare situation, and serves to highlight the importance of mechanistic modelling even as a population dynamics perspective gains importance in the cancer therapy landscape.

## 2 Methods

### 2.1 Model framework and model types

Our model describes a two cell-type cancer system, in which one cell type, *s*, is sensitive to a hypothetical drug treatment while the other, *r*, is resistant. The two cell types are assumed to grow at cell type-specific growth rates, *r*_*s*_ and *r*_*r*_ respectively within the same micro-environment, and from classical ecological theory, it follows that they inhibit each other’s growth in a density-dependent manner (May et al., 2007), with corresponding interaction strengths given *α*_*rs*_ and *α*_*sr*_. We further assume that cancer cells in the system can switch between the sensitive and resistant state at cell type-specific rates, *t*_*s*_ and *t*_*r*_. The full model that includes all these processes can be written as follows:

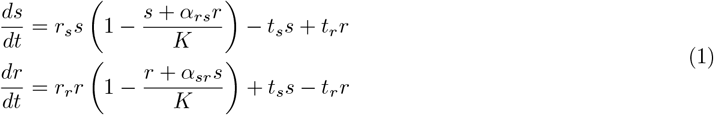

System 1 contains seven parameters: *r*_*s*_, *r*_*r*_, *α*_*rs*_, *α*_*sr*_, *t*_*s*_, *t*_*r*_, and *K*, the total carrying capacity of the system. Following previous work, we assume *K* =10,000 (Cunningham et al., 2018, Vibishan et al., 2024), resulting in six free parameters. Exploration of these parameters is discussed in a later section.

Based on their values, System 1 can be further categorised into four general model types:

- SC (symmetric competition): *t*_*s*_ = *t*_*r*_ = 0 and *α*_*rs*_ = *α*_*sr*_; *α*_*rs*_, *α*_*sr*_ *>* 0
- SC+Tr (symmetric competition + transitions): *t*_*s*_, *t*_*r*_ *>* 0 and *α*_*rs*_ = *α*_*sr*_; *α*_*rs*_, *α*_*sr*_ *>* 0
- AC (asymmetric competition): *t*_*s*_ = *t*_*r*_ = 0 and *α*_*rs*_ ≠ *α*_*sr*_; *α*_*rs*_, *α*_*sr*_ *>* 0
- AC+Tr (asymmetric competition + transitions): *t*_*s*_, *t*_*r*_ *>* 0 and *α*_*rs*_ *α*_*sr*_; *α*_*rs*_, *α*_*sr*_ *>* 0

In keeping with the stated aims of the study, we do not consider negative values of *ω*_*rs*_ and *ω*_*sr*_, which would reflect mutualistic interactions between the two cell types. The addition of mutualistic vs. competitive interactions is a complication we choose to forgo in the current study in order to keep overall model behaviour tractable.

### 2.2 Characterising steady states

Analytical solutions are known for the two-species Lotka-Volterra (LV) competition system with competitive interactions (May et al., 2007, Chapter 7), for which the fixed points are (0, 0), the two single-species points, (*K*, 0), (0, *K*), and one mixed-species point that we represent here by (*s*^*\**^, *r*^*\**^). While the origin is always a source, the stability of the remaining three points is a function of the values of *ω*_*rs*_ and *ω*_*sr*_. The theory behind this is well-known, which we shall not recount here, but (*s*^*\**^, *r*^*\**^) can be calculated explicitly in terms of *K, α* _*rs*_, and *α* _*sr*_, which will determine the relative frequencies of the two species under stable coexistence. While it is not analytically feasible to identify all fixed points of System 1, we can nevertheless understand the system behaviour in terms of what is known from the simpler LV system. To this end, we first derive conditions under which the fixed point (*s*^*\**^, *r*^*\**^) from the two-species LV model is also a fixed point of System 1. Furthermore, by analysing the behaviours of *ds/dt* and *dr/dt* in System 1 for small perturbations around this fixed point, we are able to glean insights about the stability of (*s* ^*\**^, *r* ^*\**^) when it is a fixed point for System 1. The full derivation for the existence of the fixed point in System 1 and its stability is given in Supplementary Text 1, but we note two key insights from our analysis here:

1. The fixed point (*s*^*\**^, *r*^*\**^) from the LV model is also a fixed point for the full model if 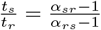.
2. For the LV model, this fixed point is stable when *α*_*rs*_, *α*_*sr*_ *<* 1 and unstable otherwise. For the full model considered here, we find that (*s*^*\**^, *r*^*\**^) is a saddle point when *α*_*rs*_, *α*_*sr*_ *<* 1 or *α*_*rs*_, *α*_*sr*_ *>* 1, and unstable otherwise. This highlights a way by which phenotypic transitions can allow for stable coexistence between two species even when inter-specific interactions are stronger than intra-specific interactions, which is generally disallowed in a system without transitions.

Given the above theoretical picture of the model’s behaviour without therapy, we use numerical simulations to gain a comprehensive understanding of the model’s dynamics in the context of a range of therapy conditions. As a first level of simplification, we categorise the overall framework into the four model types mentioned above, which allows us to analyse the effects of transitions and competition in terms of four categories rather than combinations of four continuous parameters. Furthermore, instead of dealing with the actual numbers of each cell type at steady state, we use a qualitative threshold of the fraction of resistant cells to classify the steady state. Following published literature (You et al., 2017), we classify a steady state as “Favourable” if the corresponding resistant cell fraction is less than 15%, and “Unfavourable” if it is greater than 15%. While this *ad hoc* stratification is not strictly biologically meaningful, such thresholding allows us to understand model behaviour emerging from a complex parameter space in a qualitative sense.

We characterise differences between model types primarily based on the proportion of model simulations that lead to an “Unfavourable” outcome. This is done by first fitting a generalised linear model with number of Favourable vs Unfavourable outcomes as the dependent variable and the model type as the explanatory variable, using binomial errors and a logit link function. We use the `stats` package from `R` (R Core Team, 2021, version 4.1.2) to fit the model, and further use this model to calculate odds ratios of Unfavourable outcomes for pairs of model types using appropriate functions in the `emmeans` package (Lenth, 2023, version 1.8.4-1). Wherever necessary, we calculate effect sizes using functions from the `rstatix` package (Kassambara, 2023, version 0.7.2). We also develop and use custom Python scripts to run simulations with a more specific combination of parameters for the heatmaps in Figure 1 and the time series plots in Figures 2 and 3.

**Figure 1.**
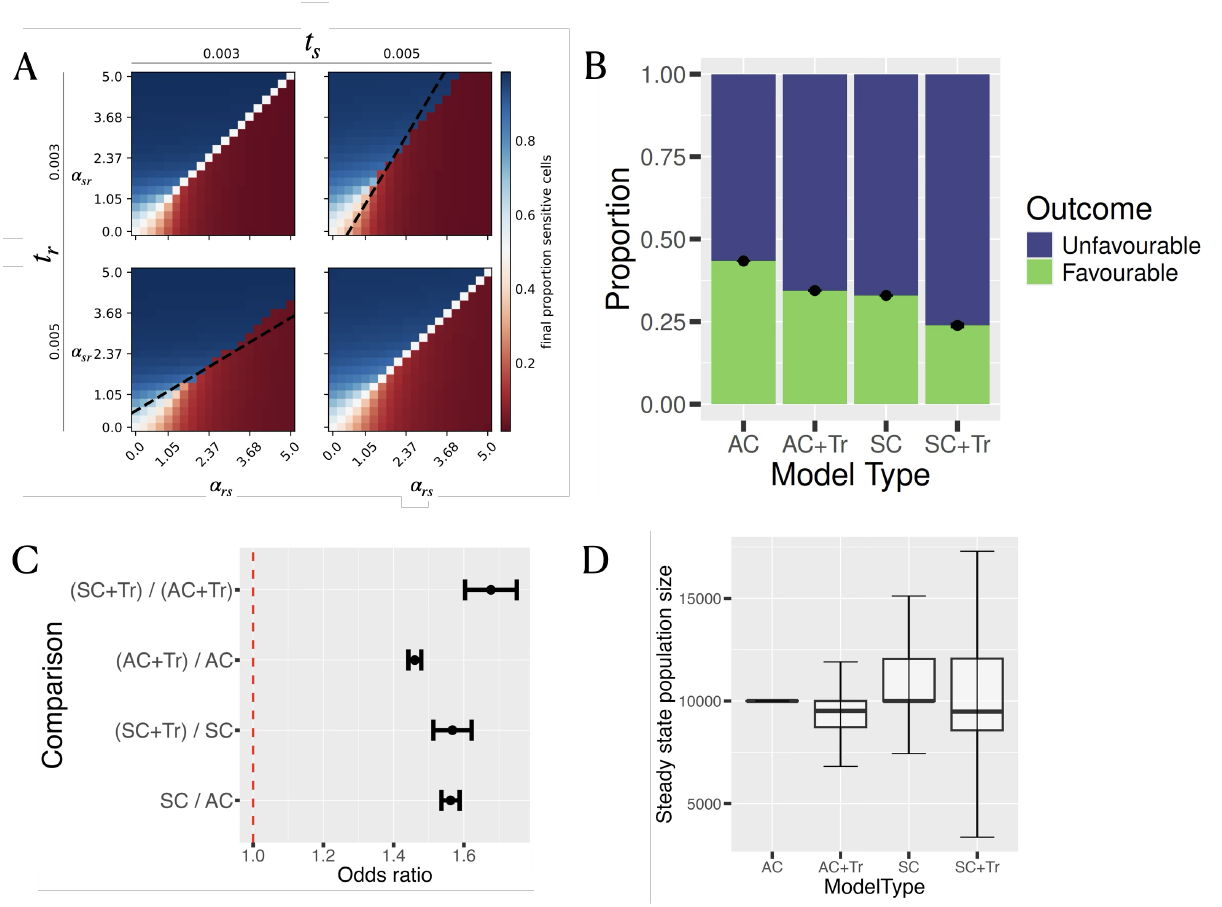
Model outcomes without therapy; (A) Steady state fraction of sensitive cells across combinations of competition and transition rates, for *r*_*s*_ = *r*_*r*_ = 0.05 and *s*(0) = *r*(0) = 100; the black dashed line is the condition 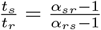 for stability of the mixed species fixed point (see the main text and Supplementary Text 1 for details); (B) Proportion of “Favourable” outcomes across model types (dots and error bars are mean ± SD over three replicate runs); (C) Odds ratio of an “Unfavourable” outcome across model types (error bars are SD over three replicate runs); (D) Steady state population size across model types (boxplots are pooled over independently-sampled parameter combinations across all three replicate runs); here and in all following figures “Favourable” implies steady state fraction of *s* ≥ 0.85. Model types, parameter definitions, ranges and sampling are described in the Methods. While both transitions and competition affect overall model outcomes, results over a large number of parameter combinations suggest that models with transitions perform worse on average in terms of “Favourable” outcomes.

**Figure 2.**
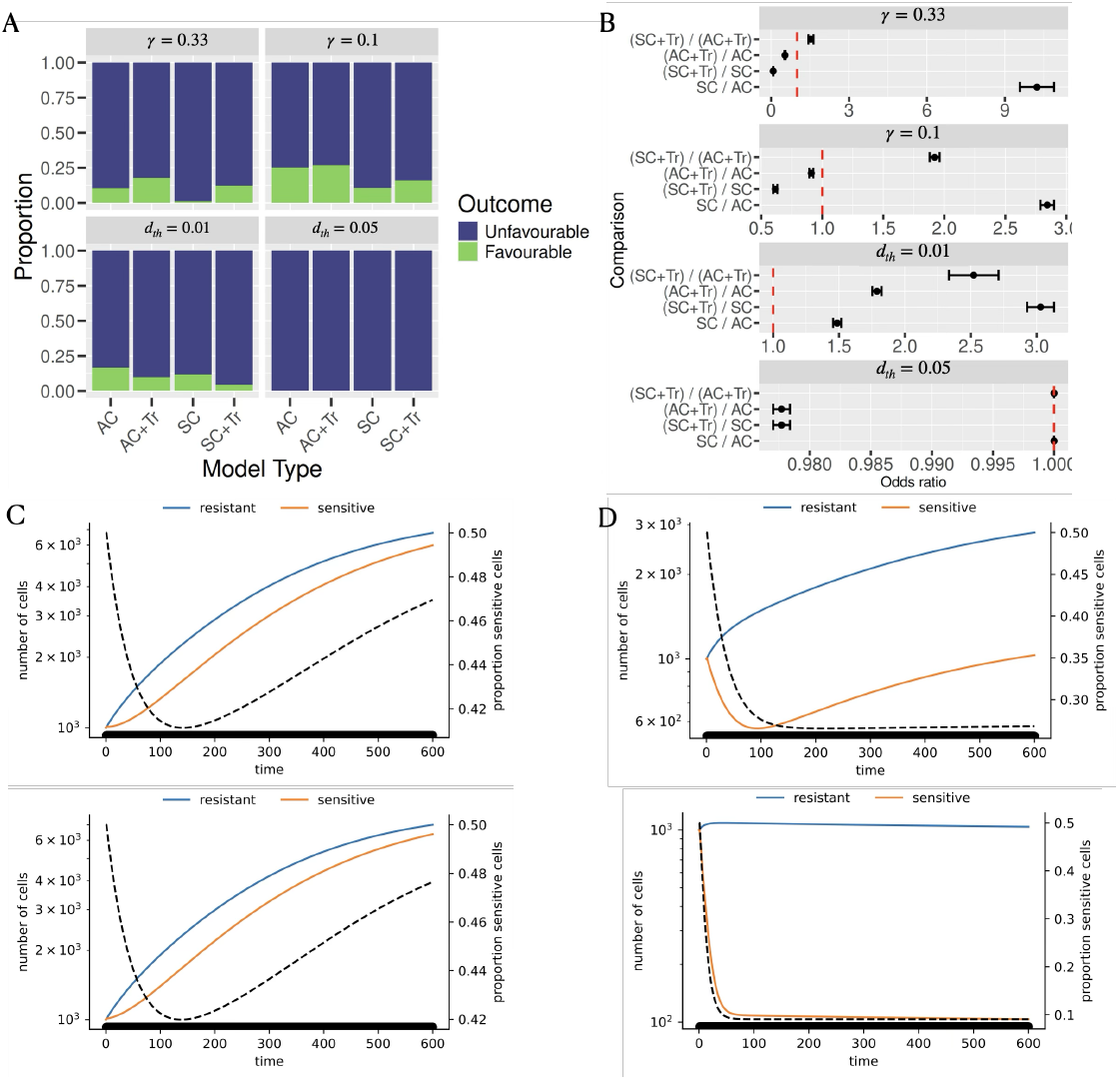
Model outcomes under constant dose therapy; (A) Proportion of “Favourable” outcomes, and (B) odds ratio of an “Unfavourable” outcome across model types. For cytostatic therapy, *ε* = 0.33 or 0.1, while for cytotoxic therapy, *d*_*th*_ = 0.01 or 0.05 (see System 2); note that the x-axis in (B) across rows is not the same. (C) and (D) Temporal dynamics of sensitive and resistant cells for two doses of (C) Cytostatic therapy (top-*γ r*_*s*_ = 0.0001, bottom-*γ r*_*s*_ = 0.001, and (D) Cytotoxic therapy (top-*d*_*th*_ = 0.025, bottom-*d*_*th*_ = 0.1. For the time series, *s*(0) = *r*(0) = 1000, *r*_*s*_ = *r*_*r*_ = 0.01, *α*_*rs*_ = *αsr* = 0.2, and *t*_*s*_ = *t*_*r*_ = 0.01; the black dashed lines show the proportion of sensitive cells, as given on the right y-axis, and the bold patches on the x-axis indicate the time points when therapy was active.

**Figure 3.**
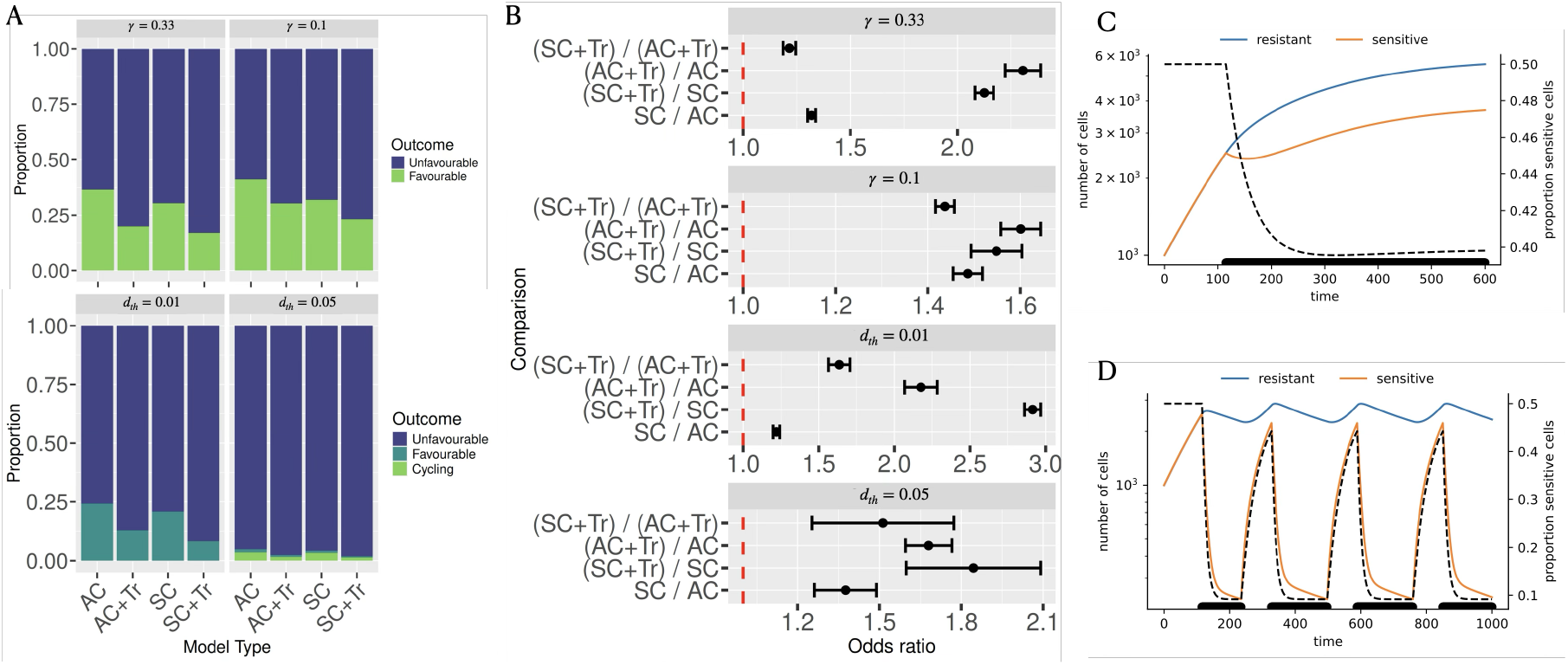
Model outcomes under adaptive therapy; (A) Proportion of “Favourable” outcomes, and (B) odds ratio of an “Unfavourable” outcome across model types. (C) and (D) Temporal dynamics of sensitive and resistant cells under cytotoxic therapy; (C) *d*_*th*_ = 0.01, (D) *d*_*th*_ = 0.1. All other definitions and parameter values are the same as previous figures.

### 2.3 Simulations with and without therapy

We simulate System 1 using Python routines from the `scipy` package (Virtanen et al., 2020). For the most part, we explore large ranges for all six free parameters as described below, except for some biologically-feasible assumptions. We assume that neither intrinsic growth rate exceeds 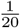 hour ^−1^based on reported literature from *in vitro* cell lines (Demicheli et al., 1991, Cowan et al., 2004), and explore values of growth rates over about two orders of magnitudes up to the assumed maximum. This also sets the timescale of the simulations in hours. Furthermore, we also assume that transition rates do not exceed the intrinsic growth rate based on known timescales of epithelial-mesenchymal transitions in some cancer types (Karacosta et al., 2019, Deshmukh et al., 2021), and explore a range of values one or two orders of magnitude below the growth rates. Both these constraints have been relaxed to a limited extent for the heatmap and time series visualisations, where transition rates of the same order of magnitude of the growth rates have been assumed occasionally. Finally, competition coefficients can be equal to, greater than, or less than one, with each scenario leading to distinct steady states. It has long been known that intraspecific competition being stronger than interspecific competition can allow for stable coexistence among species at steady state (Case, 2000, May et al., 2007). We therefore explore values of the competition coefficients over a seven-to ten-fold change both above and below one. Table 1 lists all free parameters and the corresponding ranges explored. We use a two-fold sampling strategy for all the parameters in the model. For each parameter, 5000 values are sampled using a Latin hypercube method while 5000 values are sampled from a log-uniform distribution. The ranges listed in Table 1 show the widest window for each parameter. We use log-uniform sampling particularly to ensure sufficient coverage towards the lower end of the parameter range, as Latin hypercube sampling can sometimes under-represent the extremes of a distribution when parameter values range over several orders of magnitude. Our sampling results in distributions that are largely uniform for both *r*_*s*_ and *r*_*r*_, while the transition rates and competition coefficients are sampled with a bias towards the lower end of the range.

**Table 1:**
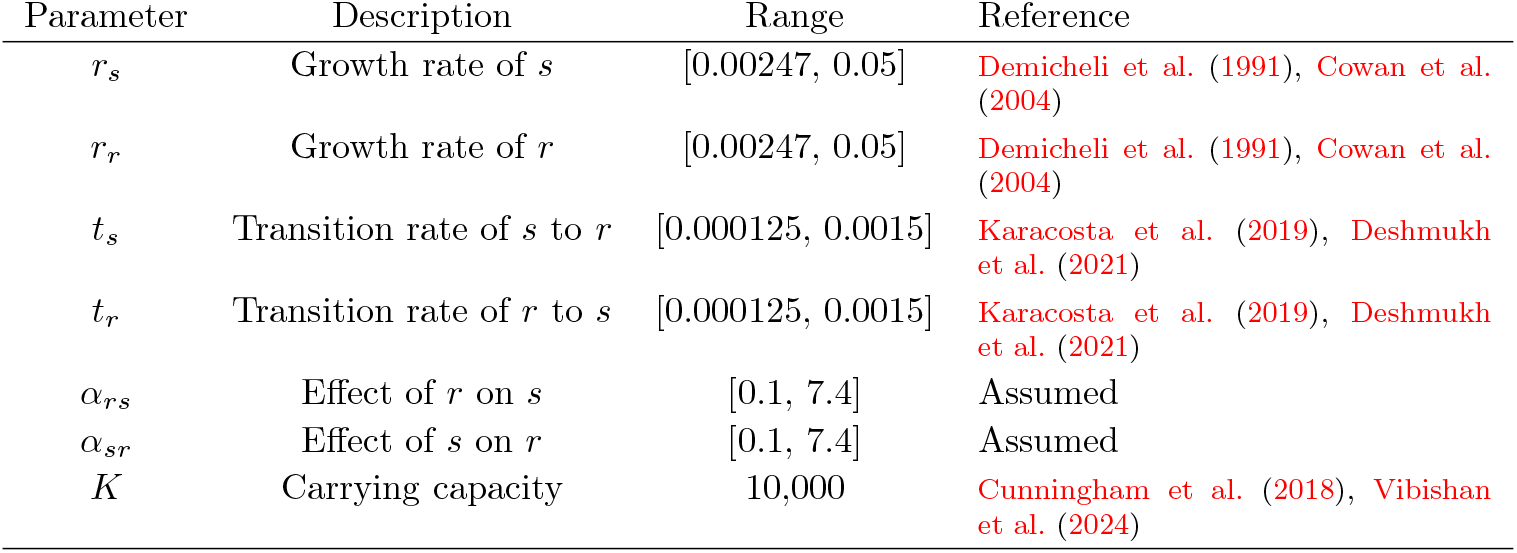
List of all parameters and their ranges explored in the model.

We run three sets of simulations in total. Each set of simulations consists of the model being run over three independent samples of parameter combinations, which serve as independent replicates of the parameter space. Each sample consists of 10,000 combinations of the six free parameters, drawn as mentioned above. The first set of simulations corresponds to the no therapy condition, and all sampled parameter values are used as is in the model. The second set corresponds to the parameter combinations for constant dose therapy, while the third set corresponds to those for adaptive therapy. Each set of 10,000 parameter combinations is further replicated across the four model types, such that between any two model types, the same sample of parameter combinations is used with appropriate modifications to the transition rates or competition coefficients. Specifically, a random sample of 10,000 parameter combinations with non-zero values for all six parameters represents a sample parameter set for the AC+Tr model. The same sample is replicated with both transition rates set to zero for the AC model, with *α*_*sr*_ set to be equal to *α*_*rs*_ for the SC+Tr model, and with transition rates set to zero and competition coefficients set to be equal for the SC model. Therefore, in each sampling of the parameter space, intrinsic growth rates for all four model types are the same, and between any two model types that differ by only one process, all other parameter values are the same. For instance, for a given sample of parameter combinations, the AC+Tr and SC+Tr models differ only in their competition coefficients while the intrinsic growth rates and transition rates of the two cell types are the same.

System 1 describes the competition and transition of drug-sensitive and -resistant cells. To capture the effect of cytostatic and cytotoxic therapy on the sensitive cells, we set

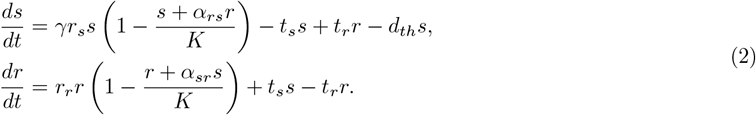

A choice of *γ* ∈ [0, 1) corresponds to cytostatic therapy, while a positive, additional death rate *d*_*th*_ *>* 0 corresponds to cytotoxic therapy. For this study, we consider *γ* = 0.33 or 0.1, corresponding to a 66.7% and 90% reduction in the normal growth rate, respectively, and *d*_*th*_ values of 0.01 and 0.05. We note that therapy does not directly affect resistant cells. Each simulation is initialised with 50 cells of each type and simulated up to 5000h, at which point steady state behaviour has long emerged. All steady state analyses mentioned above are carried out on each independent replicate, in order to obtain error estimates for odds ratios, effect sizes, and the net proportion of unfavourable outcomes.

For constant dose therapy, the drug treatment is kept constant from the beginning of the simulation regardless of the current population size. For adaptive therapy, therapy is only turned on when the total population size exceeds 50% of the carrying capacity, and turned off when the total population size drops below 10% of the carrying capacity. It is straightforward to see that while the constant dose treatment conditions would result in a clear steady state, adaptive therapy may give rise to cyclic population dynamics, which cannot meaningfully be stratified into “Favourable” and “Unfavourable”. For these cases, we use a custom `R` function to calculate the frequency spectrum of each time series and identify the frequency with the highest spectral density. The inverse of this frequency gives the time period of fluctuations in the time series. Under adaptive therapy therefore, those time series with zero frequency are assumed to be non-cycling and stratified into favourability categories based on the resistant cell fraction. Those time series with positive time periods under adaptive therapy are identified as cycling, with no further stratification based on the resistant cell fraction.

### 2.4 Virtual cohort simulations

In the therapy simulations described above, the initial frequencies of sensitive and resistant cells are all assumed to be the same as each simulation begins with 50 cells of each type, unless mentioned otherwise. However, this does not reflect the fact that in the clinic, cancer patients are diagnosed over a range of time points with considerable variability in the amount of tumour growth at the time of diagnosis (Carter et al., 1989, Rosenberg et al., 2005). More specifically, constant dose therapy (CDT) in the previous section is initiated at a tumour population size of about 100, which is much lower than the clinical detection limit for solid tumours, and therefore likely too early compared to when therapy is administered in a clinical setting (it might, however, be relevant to chemotherapy following surgical intervention with negligible residual disease). In order to account for this gap with clinical practice, we run an independent set of simulations, in which about 10,000 combinations of the six free parameters are sampled from a log-uniform distribution (same ranges as earlier; Table 1) across all four model types. We sample initial cell numbers for both cell types from a uniform distribution in the range [10, 100] and simulate the model in absence of therapy for each parameter combination for an initial period of 1000h. At the 1000h time point, we introduce cytotoxic therapy with *d*_*th*_ = 0.05, and simulate the model for a further 5000h, in order to reach a steady state. This whole exercise is carried out independently for constant dose therapy as well as adaptive therapy. It is straightforward to see that under the constant dose regime, therapy is active from the 1000h mark onwards when it is introduced, but the turning on of therapy under the adaptive regime is dependent on the total population size at the 1000h time point.

As with previous simulations, we stratify the steady state and cycling behaviour of each parameter combination into a “Favourable”, “Unfavourable”, or “Cycling” outcome, which is then compared between the 1000h time point just before therapy (BT), and the eventual steady state under either constant dose therapy (CDT) or adaptive therapy (AT). Furthermore, we are also interested in understanding if either the population size or the fraction of sensitive cells just before the introduction of therapy can predict the steady state population size and fraction of resistant cells after therapy. Accordingly, we address this question using separate principal component analyses (PCAs) for each therapy (CDT and AT) and each variable type (population size and resistant cell fraction), all normalised to the same scale. All six model parameters are also included and the first two principal components are chosen for visualisation and interpretation. We use the `prcomp` function from the stats package (R Core Team, 2021, version 4.1.2) for the PCA itself, and appropriate functions from the `ggplot2` library (Wickham, 2016, version 3.5.1) and `ggbiplot` package (Vu and Friendly, 2024, version 0.6.2) for all the figures.

### 2.5 Parameter identifiability analyses

The clinical relevance of testing specific therapy protocols using the current model framework is self-explanatory, since these protocols are directly related to those currently being considered for a range of clinical applications. However, a broader theoretical question also arises concerning the ability to identify specific parameter values corresponding to specific temporal behaviours in the model. While theoretical studies can use simulations to associate model behaviours with pre-determined parameter values, the inverse problem of inferring model parameters based on actual time course data can be non-trivial. In particular, the structure of the model can circumscribe the unique identification of its parameters, and this limit can be understood in terms of the structural identifiability of a model. Here, we perform structural identifiability of System 1 categorised into four general model types using the GenSSI package (Ligon et al., 2018). The package uses a Lie derivative-based approach to convert ODEs to a polynomial system that can provide useful information regarding mathematical relationships between free parameters in the model, and how this determines which parameter values, (1) can be identified independently of other parameters (globally identifiable), (2) can be identified but depending on other parameters (locally identifiable), or (3) cannot be identified (non-identifiable), given information of one or more model output variables. We test the structural identifiability of System 1, assuming one or more time series of the total population size with known/unknown fractions of *s* and *r* cells in the population at time, *t* = 0 are available.

## 3 Results

This study aims to investigate the influence of phenotypic plasticity as well as density-dependent competition on the long-term dynamics of a heterogeneous population of sensitive and resistant cancer cells. We use a classical Lotka-Volterra (LV) competition model framework with modifications that describe switching in cell phenotype between the sensitive and resistant cell states, and investigate the implications of such a model’s behaviour for therapy design. The primary response variable for our model framework is the fraction of resistant cells at the end of each simulation. This fraction is used to classify the outcome of each simulation as either Favourable or Unfavourable, as mentioned earlier. We study how this fraction varies as a function of each individual parameter in the model to obtain a qualitative indication of the roles played by each of the parameters. We consider 6 independent free parameters – intrinsic growth rates of sensitive and resistance cell types, competition coefficients that describe the density-dependent effect of one cell type on the other, and rates of phenotypic transitions between the two cell types. As mentioned earlier, combinations of competition and transition parameters give rise to four general model types – symmetric vs. asymmetric competition, with vs. without transitions – and we therefore examine model outcomes in terms of these four model types.

### 3.1 Transitions and competition both affect model outcomes in the absence of therapy

Figure 1A displays proportion of cells that are sensitive in the long-term for varying combinations of the competition and transition rates. The trends largely conform to the expected stability behaviour of the mixed-species fixed point of the LV model. The parameter combinations on the main diagonal in both the top left and bottom right panels correspond to 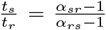 and the mixed-species fixed point is, as expected from LV theory, stable in this parameter region. The black dashed line represents the same constraint in the other two panels, when the transition rates are not equal. The mixed-species fixed point is clearly stable for a region around the line when inter-species competition is low (*α*_*rs*_, *α*_*sr*_ *<* 1), but resolves into a single-species equilibrium if inter-species competition is high. Together, this reveals the complex interplay between competition and transition rates that controls the overall steady-state behaviour of the model.

Figure 1B shows a slightly different perspective of model behaviour that considers the overall effects of competition and transitions as processes over a large number of parameter combinations. Across model types, more than half the simulations resulted in an Unfavourable steady state, implying a resistant fraction greater than or equal to 15%. The propensity to be classified as Unfavourable also varies across model types. An odds ratio analysis reveals that models with non-zero transition rates are on average more likely to result in an Unfavourable outcome than those with no transition terms (Figure 1C). Likewise, symmetric competition models are also associated with higher odds of an Unfavourable outcome than asymmetric competition models, with or without transitions.

The long-term size of the cancer cell population also depends on the model type (Figure 1D). Models with either symmetric competition or plasticity or both tend to show greater variability in the steady state population size. Figure 1D also suggests that for a given type of competition, models with transitions tend to have a lower steady state population size, whereas models with symmetric competition tend to have a higher steady state population size than those with asymmetric competition (see Supplementary Table 1 for the statistical comparisons).

### 3.2 Favourability of cytostatic vs cytotoxic constant dose therapies

The proportion of favourable model outcomes is markedly lower under constant dose therapy regimes than in the absence of therapy (Figure 2A, Supplementary Figure S1A). This is true for both cytotoxic as well as cytostatic interventions. The proportion of favourable outcomes decreases for the higher dose of constant dose cytotoxic therapy, while the fraction of favourable outcomes improves marginally for a higher dose of constant dose cytostatic therapy (Supplementary Table 3). This suggests that successive increases in drug dose could have different dynamical consequences depending on the mode of action of the drug. This is also clearly reflected in the temporal dynamics in Figures 2C and 2D. It is worth mentioning here that our definitions of Favourable and Unfavourable do not include the total population size, and therefore, a Favourable steady state can still be lethal in the clinical sense due to an untenably large tumour load. The lower prevalence of Favourable steady states under therapy, as seen in Figure 2 should not therefore taken to be an argument to abandon all therapy altogether.

Constant dose cytostatic therapy could potentially further distinguish the effects of competition asymmetry and phenotypic transitions, as reflected in the odds of an Unfavourable outcome across model types. Models with transitions had lower odds of an Unfavourable outcome than those without transitions, regardless of the competition type (Figure 2B). This indicates that the effect of phenotypic transitions on model outcomes has been reversed by the introduction of constant dose cytostatic therapy. On the other hand, the effect of competition asymmetry on model outcomes remains the same as under no therapy conditions, as models with symmetric competition have greater odds of an Unfavourable outcome.

On the other hand, constant dose cytotoxic therapy shows similar trends as in the absence of therapy in terms of whether transitions improve the odds of an Unfavourable outcome. For low doses of cytotoxic therapy, the presence of phenotypic transitions leads to worse odds of an Unfavourable outcome (Figure 2A,B). Likewise, models with symmetric competition are associated with worse endpoints than those with asymmetric competition.

### 3.3 Stable population cycles under adaptive therapy are rare

The main goal of adaptive therapy is to achieve stable population cycles through intermittent withdrawal of therapy, thus controlling overall tumour growth dynamics. This is observed in a very small proportion of the parameter combinations tested in this study, suggesting that this is a relatively rare occurrence in our model (Figure 3A). More importantly, we find that stable population cycles are observed only for the higher dose of cytotoxic adaptive therapy, while none of the remaining therapeutic conditions are able to sustain stable cycles (Figure 3A). This effect of higher dose is also reflected between Figures 3C and 3D. This suggests that some minimum reduction of the sensitive cell population is necessary in each round of therapy for effective implementation of adaptive therapy. Notwithstanding the rarity of stable population cycles, the number of parameter combinations leading to a Favourable outcome is considerably higher under adaptive therapy than under constant dose therapy (Supplementary Figure S1). While we cannot rule that the relative rarity of stable cycles under adaptive therapy is an artifact of our sampling strategy, within the constraints of our sampling, we are still able to identify general features of some parameters that are clearly associated with stable population cycles, which we detail below.

Apart from parameter combinations that result in stable cycles, the remaining parameter space can still be divided into Favourable and Unfavourable outcomes, as with the previous two cases. Both transitions and symmetric competition affect the odds of an Unfavourable outcome in the same way in that they both increase the proportion of endpoints classified as Unfavourable (Figure 3B). It is worth noting that presence vs absence of transitions seems to affect the odds of an Unfavourable outcome more strongly than the symmetry of competition (Figure 3B). Nevertheless, the similarity in overall trends in the odds ratios between Figures 2 and 3 suggests that the dynamical effects of phenotypic transitions and competition type on the steady state outcome remain qualitatively the same under adaptive therapy as under CDT.

### 3.4 Overall predictors of stable cycles under adaptive therapy

Given that population cycles are an endpoint of particular interest, we examine these parameter combinations more closely to identify which conditions are conducive for adaptive therapy. Figure 4A clearly shows that the time period of population cycles could be a predictor of model type, as models with transitions have a significantly higher period of oscillations than models without transitions. To further elucidate the parameters that contribute to population cycles, we evaluated the effect size between parameter values associated with cycling vs non-cycling dynamics. Figure 4C demonstrates that relatively lower transition rates, stronger competition (particularly for the effect of sensitive cells on resistant), and low intrinsic growth rates of the resistant cell type constitute conditions that lead to population cycles. In particular, the parameter *α*_*sr*_ that describes the density-dependent effect of sensitive cells on resistant cell growth has the highest effect sizes across model types (Figure 4B), which indicates that this parameter is a strong determinant of the occurrence of stable population cycles.

**Figure 4.**
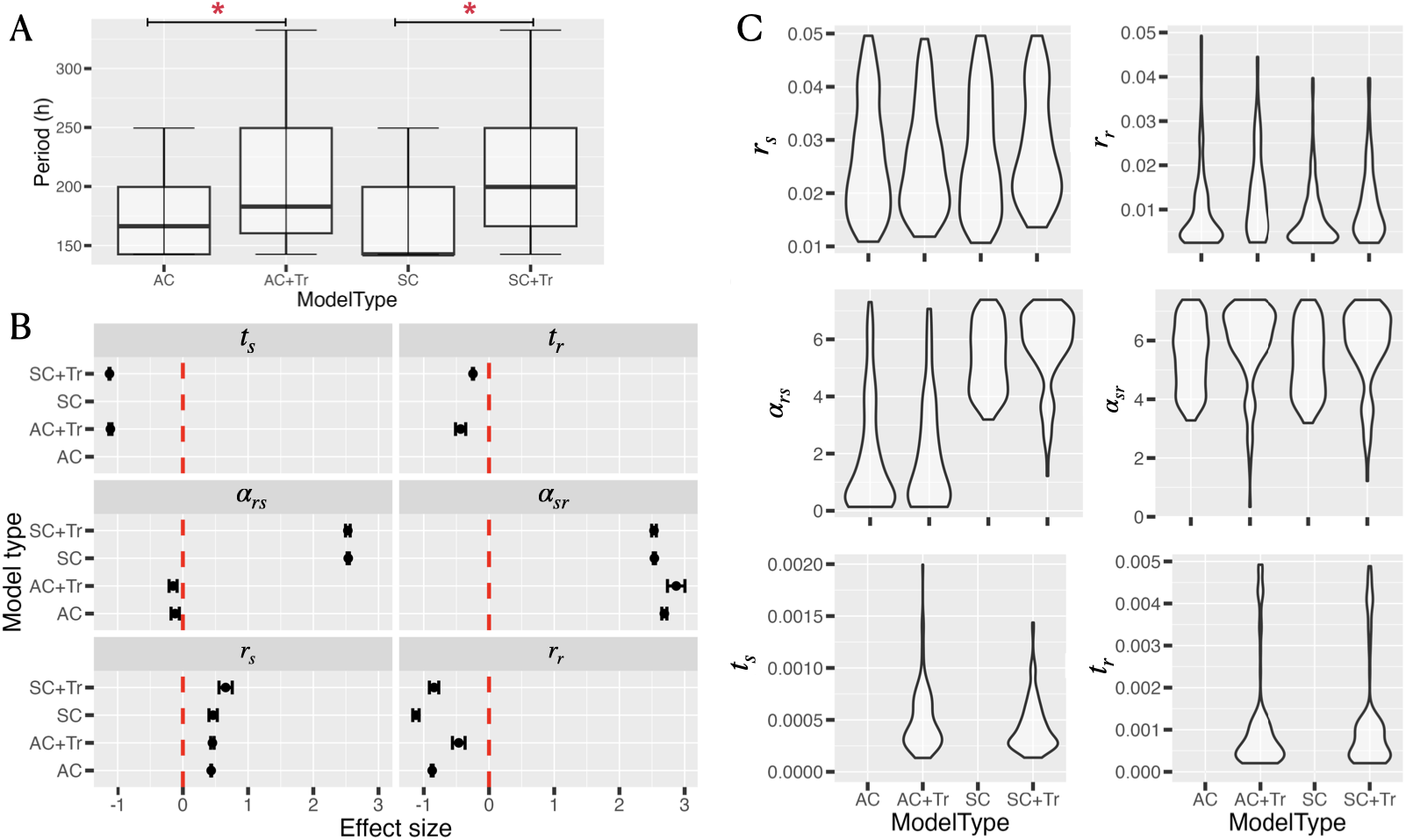
Model conditions associated with stable population cycles; (A) Distribution of the periodicity of oscillations across model types. Statistical significance was assessed using the Wilcoxon signed-rank test (see Supplementary Table 3); (B) Effect size of the parameter distributions with cycling vs. non-cycling outcomes, quantified using Cohen’s *d*. Positive effect sizes indicate larger respective parameter values in cycling outcomes; (C) Distributions of all parameter values that resulted in population cycles across model types.

It is worth pointing out that while *α*_*sr*_ is large in all models with stable population cycles, *α*_*rs*_, which describes the density-dependent effect of resistant cells on sensitive cell growth, must be particularly weak in models of asymmetric competition in order to achieve stable cycles. This suggests that under conditions of asymmetric competition, the combination of large *ω*_*sr*_ and small *ω*_*rs*_, which essentially produces a balance in competition strongly favouring the sensitive cell type, is particularly suited for stable population cycles under adaptive therapy. Interestingly, the general effect of competition asymmetry is not affected by the presence of phenotypic transitions, suggesting that in this case–for these transition rates and competition strengths–competition plays a stronger role in determining population dynamics under adaptive therapy.

### 3.5 Delayed therapy in virtual cohorts favours adaptive therapy

While the results so far illustrate therapeutic response as a function of dynamical processes within the tumour population, the number and frequency of sensitive vs. resistant cells is also likely to determine how the tumour population responds to a given treatment. Our virtual cohort simulations address this by delaying the onset of therapy by a fixed time period during which growth may occur. This delay could in turn lead to therapy being applied to a very different composition of cells compared to the simulations so far. Figure 5A illustrates the marked effects of population composition. On average, constant dose cytotoxic therapy has almost no Favourable steady state outcomes across model types, while the number of Favourable outcomes under adaptive therapy is clearly much smaller than the previous case (Figure 3). PCAs reveal no consistent associations of either resistant cell fraction or the steady state population size with any of the model parameters (Figure 5B and C), for either CDT or AT. This belies the complexity of interactions between the parameters and model outcomes, with no single parameter showing an overwhelmingly strong effect. On the whole however, the delay in treatment onset has a clear negative effect on the treatment outcome, and models with transitions are still consistently at a higher odds of an Unfavourable outcome.

**Figure 5.**
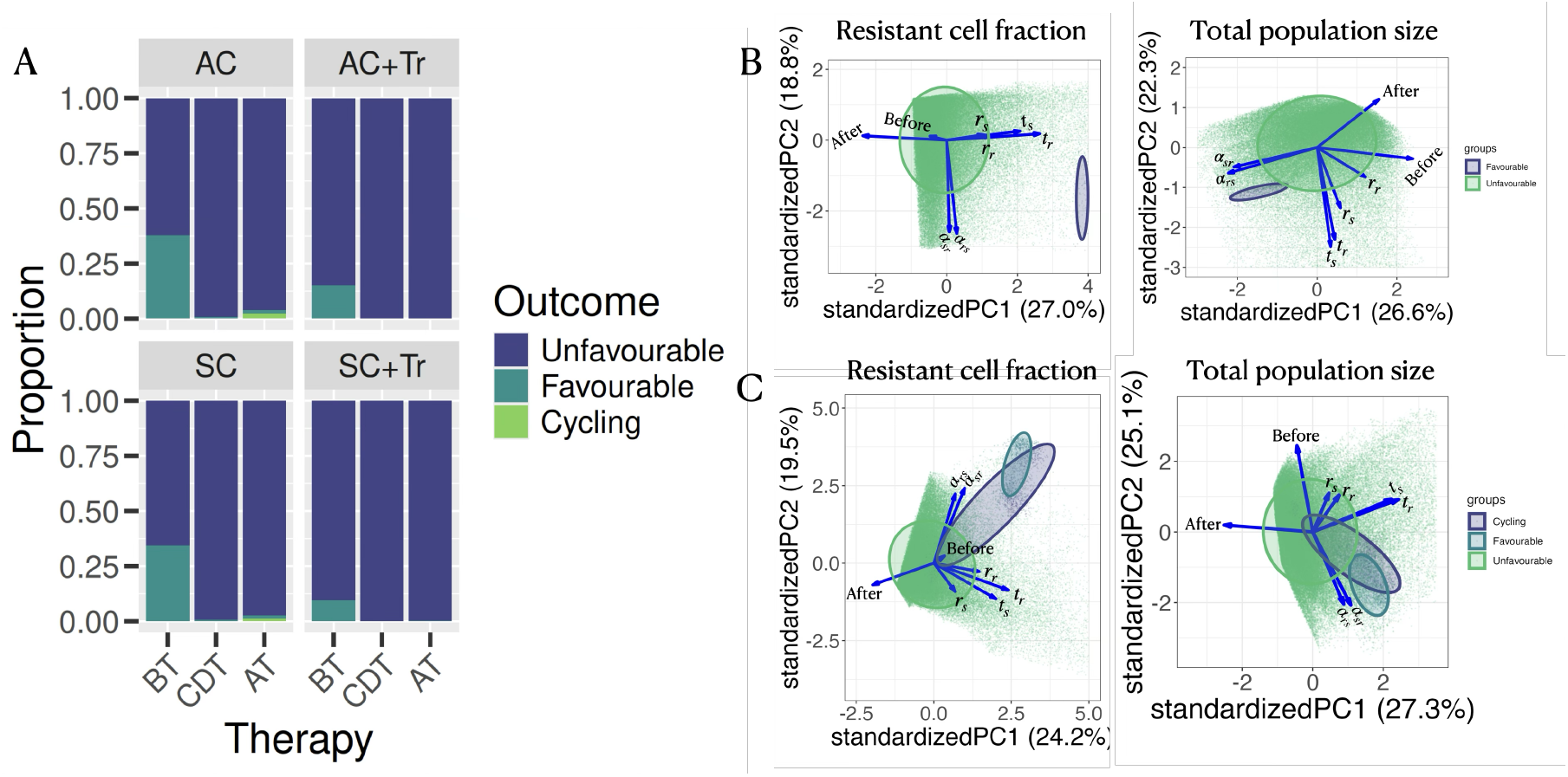
Therapy outcomes for the simulated virtual cohort with cytotoxic therapy (*d*_*th*_ = 0.05); (A) Proportion of “Favourable” outcomes across model types before therapy onset (BT), and at steady state after constant dose therapy (CDT) or that after adaptive therapy (AT); (B) and (C) Biplots of PC1 against PC2 for (B) constant dose therapy (CDT) and (C) adaptive therapy (AT). “Before” and “After” refer to the values of the parameter of interest as indicated above each panel before the onset of therapy and at steady state with therapy respectively.

### 3.6 Structural identifiability of parameters

Figure 6 shows the results of identifiability analyses for all model parameters in System 1. This shows that when the only available data is a time series of the total cell count (*s*+*r*), then regardless of whether the relative frequencies of the sensitive and resistant cells are known/unknown to begin with, none of the parameters is identifiable for System 1. However, the identifiability of the model parameters, and consequently the model, improves when two or more time series are available from different initial conditions. Taken together, this indicates that the mathematical structure of this general class of ODE-based Lotka-Volterra models imposes certain relationships between the parameters that make it particularly difficult to map a given temporal behaviour to a unique set of parameter values. Detailed results of parameter-wise identifiability for all models can be found in the Supplementary Text.

**Figure 6.**
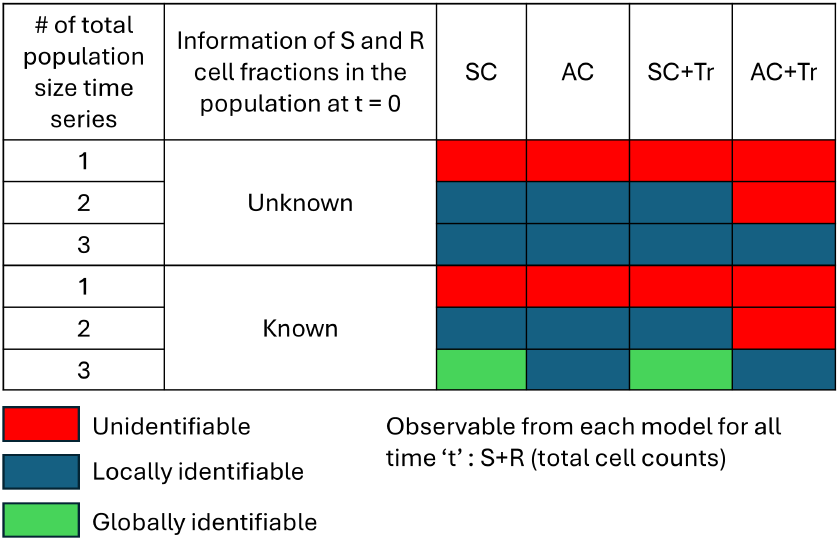
Results of the structural identifiability analyses. For this analysis, we consider total cell count as the observable variable at all times and estimate of sensitive and resistant cell fractions at the initial time point is either unknown/known. Globally identifiable signifies that the model parameters can be estimated uniquely, Locally identifiable signifies that the model parameter’s values depend on the values of other globally or locally identifiable parameters of the model, and Non-identifiable signifies that model parameters cannot be estimated for the given observable variable.

## 4 Discussion

Adaptive therapy essentially attempts to use tumour-internal processes to control the overall growth of the tumour. In this context, the results from this study highlight the importance of phenotypic plasticity both for the effective implementation of novel therapeutic strategies as well as the scope of dynamical modelling in a clinical setting.

Our simulation results show that phenotypic plasticity, or the ability of cells to switch between sensitive and resistant states, is broadly detrimental to effective therapy using a cytotoxic drug (Figures 2B and 3B). Likewise, our results also identify symmetric competition between sensitive and resistant cell types to be further detrimental to therapeutic outcomes across model types and therapy modalities. Interestingly, specifically under constant dose cytostatic therapy, we find that the presence of phenotypic plasticity is associated with a better therapeutic outcome. This could be due to the fact that while a cytostatic drug reduces the growth rate of sensitive cells, it does not remove them from the population instantly, which could help maintain the biotic inhibition exerted by sensitive cells on resistant cells. However, this line of reasoning does not explain the observed trends under cytostatic adaptive therapy: here, the presence of plasticity leads to worse outcomes as was the case without therapy. This is surprising, as it is straightforward to expect such dynamic shifts between a resistant-dominant and sensitive-dominant population to occur easily during drug holidays. Nevertheless, we identify at least three conditions for efficacious therapy design: (1) cytostatic drugs are better applied in a continuous manner, (2) the extent of phenotypic plasticity must be minimised, and (3) asymmetry in the competitive interactions between sensitive and resistant cells must be encouraged. Needless to say, these three conditions are not equally feasible to implement. While well-characterised cytostatic drugs are available for a wide range of cancers Galmarini et al. (2012), it is less clear if drugs exist that can tune the dynamics of intra-cellular processes that lead to phenotypic plasticity (but see Liu et al., 2012). The exact basis of this plasticity is still an area of active investigation Caiado et al. (2016), and may even vary across cancer types. Directly affecting the competition between resistant and sensitive cells is a tricky proposition, but recent work is beginning to highlight instances of differential nutrient use between resistant and sensitive cell types (Immanuel et al., 2021, Kareva and Brown, 2021, Emond et al., 2023, Vibishan et al., 2024), and connections between metabolic states and drug resistance have long since been actively investigated (Massie et al., 2011, Vaz et al., 2012, Archetti et al., 2015). Taken together, this indicates that a mechanistic understanding of tumour-internal dynamics could lead to strategies that could attenuate the balance of competition within the tumour to improve therapeutic outcomes. Here, it is worth noting that these speculations are underlined by a shortcoming of this study in its current form – with most of our results relying on steady state outcomes, definitive comments on the underlying temporal dynamics leading to these outcomes are not possible. Analyses that are more focused on identifying shifts in the balance between transition and density-dependent growth over the course of a simulation could help to shed more light on the temporal details of how these trends emerge.

Our analyses of therapy administration with the simulated virtual cohort highlight the importance of the tumour composition at the point of therapeutic intervention in determining the eventual outcome. When starting with 50 sensitive and 50 resistant cells, constant dose therapy has generally poorer outcomes than adaptive therapy (Supplementary Figure S1). This remains the same when therapy is introduced after a phase of tumour growth, but overall treatment outcomes with either CDT or AT are much worse when administered following a period of no-therapy growth. This suggests that the ability of constant dose or adaptive therapy to control or affect population growth strongly depends on the initial state of the population, but indicates that adaptive therapy may be slightly better with tumours diagnosed at a later stage.

In the same vein, a recent study with a Lotka-Volterra system similar to the current model but without phenotypic plasticity between sensitive and resistant cells presents some relevant findings (McGehee and Mori, 2024). The model compares adaptive therapy through two modalities – intermittent, which is similar to the modality implemented in the current study and involves dose skipping, and constant dose, which involves continuous modulation of the drug dose without skipping. In the first place, it is interesting to note that despite the presence of phenotypic plasticity in the current study, both their model and ours identify large parts of the parameter space for which adaptive therapy by dose skipping is not a viable therapeutic strategy. However, McGehee and Mori (2024) find that when the sensitive cells are strong competitors, constant dose “anti-proliferative” treatment, which we have termed cytotoxic in the current study, outperforms any kind of adaptive dosing regime. This is broadly consistent with our results reported here, and a comparison of intermittent (i.e., on-off type) dosing regimes against constant dose adaptive regimes with dose modulation could be a useful extension of the current work, particularly given that McGehee and Mori (2024) suggest the latter could induce less drug-related toxicity. However McGehee and Mori (2024) also assume a cost of resistance in all of their results, which is not part of the current study. The exact role of a cost of resistance therefore requires further investigation.

The results from the numerical simulations presented here provide a broader perspective than what is reflected in the analytical results alone (Supplementary Text 1). The mathematical conditions for stability of co-existence of sensitive and resistant cells reveal an unexpected way by which transitions enable two-species co-existence at steady state even when inter-specific interactions are stronger than intra-specific interactions. In the context of theoretical ecology, this is an interesting finding, and parallels the well-known dispersal-competition tradeoff that can allow co-existence of two competing species in patchy habitats (Amarasekare and Nisbet, 2001). In the context of cancer however, these results lack an ensemble-level view of the model behaviour over the entire parameter space that is more effectively captured by the larger-scale numerical simulations.

There are a few ways in which observations from this study are directly relevant to the clinical implementation of adaptive therapy. In the ideal case, adaptive therapy could be designed accurately if the current frequency of sensitive cells is known at any given time point. However, data of such granularity are rarely available for real tumours, and therapy design usually relies on some marker of total tumour volume (Djavan et al., 2012, Brady-Nicholls et al., 2020). The effectiveness of such markers is naturally limited by the fact that by themselves, they are uninformative about underlying system processes (Figure 6). Our data show that in cases where therapy leads to stable population cycles, the time period of fluctuations could be related to the nature of phenotypic plasticity in the system. Specifically, we find that the presence of phenotypic plasticity leads to a longer time period of oscillations when therapeutic conditions lead to stable population cycles (Figure 4A). In the same context, we also find that strong competition from the sensitive cells is a necessary and sufficient condition for stable population cycles. Such observations are ripe for further investigation as it suggests that (1) features of the underlying system could be reflected in the dynamics of the total population size even without additional biomarkers of specific sub-populations in the tumour, and (2) some model output (in this case, time period of fluctuations), along with qualitative knowledge of the underlying tumour biology (in this case, whether or not sensitive cells are strong competitors), could be used to infer which processes are active within the tumour.

The other major clinical implication of this modelling framework concerns the identifiability of parameter values. As mentioned above, clinical data are generally sparse in terms of detailed cell-level or micro-environmental information, and therapy design and implementation is usually based on broad indicators of tumour growth. With such sparsity, it remains unclear if phenomenological model frameworks of the kind used in this, and many other studies, can be satisfactorily parameterised in order to be useful in personalised therapy design. In fact, our results indicate that global identifiability of parameters in such a phenomenological model framework requires not only multiple time series data, but also multiple initial conditions of the relative frequencies of sensitive and resistant cells–conditions that are highly unlikely to be met in a clinical setting. This practical lack of parameter identifiability places serious constraints on the extent to which such models can be applied to design and implement therapy, particularly given that the parameters most relevant to tumour-internal processes (transition, competition, etc.) cannot be reliably determined under realistic assumptions of data availability. Since this is a structural limitation of this class of models, solutions to this will need either further careful experimental estimation for some parameters, or alternative modelling frameworks that might be compatible with existing clinical data while also being aligned to emerging sources of data from tumours. Whether these are deep-learning/AI-based solutions, or more conventional mechanistic modelling approaches, or some combination thereof, will only become clear in the coming years.

While no model can account for every possible factor of importance, some key features are nevertheless worth highlighting that we have not considered in the current study. First, the rates of phenotypic transitions in our model are constant, regardless of the presence of drug. This could be an important limitation, particularly given recent theoretical work, which has demonstrated that drug-induced plasticity could reduce the efficacy of constant dosing regimes. Instead, pulsed dosing regimes are seen to perform in delaying the onset of long-lasting drug resistance in the population (Gunnarsson et al., 2024, Gevertz et al., 2024). On the other hand, other models that account for drug-induced plasticity have shown that long-term resistance acquisition is favoured primarily when the characteristic timescale of phenotype switching is much larger than that of cell growth (Gunnarsson et al., 2020). This reflects the parameterisation of our model, as well as the fact that the parameter space leading to viable therapeutic outcomes in our results is quite marginal. Nevertheless, this can be considered circumstantial evidence at best – future modelling efforts must be modified to include drug-induced plasticity in order to corroborate any expected impacts of the timescales of plasticity vs cell division on overall population dynamics. On a related note, cellular memory is known to play a crucial role in determining the speed and accuracy of adaptation in biological populations under fluctuating environmental regimes (George, 2023). Such dynamical effects are naturally highly pertinent to adaptive therapy, which is inherently fluctuating. The exploration of memory-driven responses to adaptive therapy would therefore be a valuable extension of the current work. Finally, all our results in this study assume a well-mixed domain, and therefore neglect any effects of spatial structure in cell-cell interactions on therapy response. Again, the importance of spatial structure in determining therapeutic success has been well-established (Bacevic et al., 2017, Strobl et al., 2022), and extensions of the current work to spatially-explicit contexts is a necessary next step in making the findings of the current model broadly relevant to cancer treatment and management.

## Acknowledgements

The authors are grateful to all members of the Cancer Systems Biology group and Tomas Gedeon for useful discussion and critical feedback.

## Author contributions

MKJ and JTG supervised research and acquired funding. MKJ, JTG, PJ, KH and BV conceptualised research. BV, PJ, VS and KH performed research. BV, PJ and CK contributed new methodology and visualisation aspects. BV wrote the original draft. All authors contributed to data analysis, and reviewing and editing the manuscript.

