## Supplementary Text 1 for "Impacts of competition and phenotypic plasticity on the viability of adaptive therapy"

Stability analyses and fixed point conditions

The current model as given in System 1 is an extension of the two-species Lotka-Volterra (LV) competition model, for which the fixed points and stability conditions are well-known (Case, 2000, Chapter 14). In the first two sections below, we derive conditions in terms of parameter values or ranges for which the fixed points from the simpler two-species LV model are also fixed points for the extended model presented in this study. We also derive criteria that provide some indications of the stability of these fixed points in the extended model. In the last section, we develop an alternative approach to investigate the steady state behaviour of the model using the geometry of conic sections.

### S1 Applying the fixed points of the two-species LV model to the extended model

The two-species LV model can be written as follows:

$$\begin{aligned} \dot{s} &= r_s s \left(1 - \frac{s + \alpha_{rs} r}{K}\right) =: f_s(s,r), \\ \dot{r} &= r_r r \left(1 - \frac{r + \alpha_{sr} s}{K}\right) =: f_r(s,r). \end{aligned} \tag{S1}$$

The fixed points can be found by setting $\dot{s} = 0$ and $\dot{r} = 0$. It is straightforward to see that $(0,0)$, $(0,K)$ and $(K,0)$ are all fixed points, while one co-existence equilibrium is also possible, which we denote by $(s^*,\ r^*)$. This can be found from the equations of the two nullclines, $s + \alpha_{rs} r = K$ and $r + \alpha_{sr} s = K$, which yields the following coexistence fixed point:

$$(s^*,\ r^*) = \left[\frac{K(\alpha_{rs} - 1)}{\alpha_{rs}\alpha_{sr} - 1},\ \frac{K(\alpha_{sr} - 1)}{\alpha_{rs}\alpha_{sr} - 1}\right] \tag{S2}$$

When $\alpha_{rs} = \alpha_{sr}$, we have $s^* = r^*$, implying that the ecological niche, whose size is quantified by the carrying capacity $K$, is split equally between sensitive and resistant cells. Using linearisation around this fixed point, it can be shown that the co-existence fixed point is only stable when $\alpha_{rs} < 1$ and $\alpha_{sr} < 1$. Otherwise (i.e., when $\alpha_{rs} > 1$ and/or $\alpha_{sr} > 1$), it is unstable (Case, 2000, Chapter 14).

The extended model considered in this study includes cell state transitions, as given below:

$$\begin{aligned} \dot{s} &= r_s s \left(1 - \frac{s + \alpha_{rs} r}{K}\right) - t_s s + t_r r = f_s(s,r) - t_s s + t_r r, \\ \dot{r} &= r_r r \left(1 - \frac{r + \alpha_{sr} s}{K}\right) + t_s s - t_r r = f_r(s,r) + t_s s - t_r r. \end{aligned} \tag{S3}$$

It is straightforward to see that the LV fixed points $(0,\ 0)$, $(K,\ 0)$ and $(0,\ K)$ are all fixed points for the extended model as well. While $(0,\ 0)$ is always unstable, the stability of the two single-species fixed points depends on the co-existence fixed point. Unlike the simpler System S1, the nullclines of System S3 are not linear, making it challenging to identify this intersection. One approach to demonstrate existence and provide conditions under which the coexistence fixed point, given by Equation S2, of System S1 is also a fixed point of the extended System S3. At the fixed point $(s^*,\ r^*)$, we have, since $f_s(s^*,\ r^*) = 0$,

$$\dot{s}|_{(s^*,\ r^*)} = f_s(s^*,\ r^*) - t_s s^* + t_r r^* = -t_s s^* + t_r r^*. \tag{S4}$$

Thus, for $(s^*,\ r^*)$ to be a fixed point System S3, we require $t_s s^* = t_r r^*$.

Substituting from Equation S2, we get

$$\begin{aligned} t_s \frac{K(\alpha_{rs} - 1)}{\alpha_{rs}\alpha_{sr} - 1} &= t_r \frac{K(\alpha_{sr} - 1)}{\alpha_{rs}\alpha_{sr} - 1} \\ \frac{t_s}{t_r} &= \frac{K(\alpha_{sr} - 1)}{\alpha_{rs}\alpha_{sr} - 1}\ \frac{\alpha_{rs}\alpha_{sr} - 1}{K(\alpha_{sr} - 1)} \\ \frac{t_s}{t_r} &= \frac{\alpha_{sr} - 1}{\alpha_{rs} - 1} \end{aligned} \tag{S5}$$

This indicates that $(s^*,\ r^*)$, as in Equation S2, will be a fixed point of System S3 as well if Equation S5 is satisfied.

#### S2 Stability of the fixed point, $(s^*,\ r^*)$

To understand the stability of the fixed point, consider a small perturbation from the fixed point, given by $(s^* + \delta s,\ r^* + \delta r)$. The fixed point can considered stable if $\dot{s}|_{(s^*+\delta s,\ r^*+\delta r)}$ is positive for negative perturbations and negative for positive perturbations. These perturbations can naturally be either both in the same direction or in opposite directions, defining two distinct axes along which stability can be investigated. Assuming a general case of some small perturbation $(\delta s,\ \delta r)$ from the fixed point and abbreviating $(s^*,\ r^*)$ to $(s,\ r)$, we can calculate $\dot{s}|_{(s^*+\delta s,\ r^*+\delta r)}$ as follows:

$$\begin{aligned}\dot{s}|_{(s+\delta s,\ r+\delta r)} &= r_s(s+\delta s)\left[1 - \frac{s+\delta s+\alpha_{rs}r+\alpha_{rs}\delta r}{K}\right] - t_s(s+\delta s) + t_r(r+\delta r)\\ &= r_s(s+\delta s)\left[1-\frac{s+\alpha_{rs}r}{K}\right] - t_s s\ +\ t_r r\ -\ \frac{r_s(s+\delta s)(\delta s+\alpha_{rs}\delta r)}{K}\ - t_s\delta s\ +\ t_r\delta r\end{aligned}$$

For the fixed point $(s,\ r)$, as given by Equation S2, $1 - \dfrac{s+\alpha_{rs}r}{K} = 0$, and from Equation S4, we also have $-t_s s + t_r r = 0$. Therefore,

$$\begin{aligned}\dot{s}|_{(s+\delta s,\ r+\delta r)} &= \frac{-r_s(s+\delta s)(\delta s+\alpha_{rs}\delta r)}{K} - t_s\delta s + t_r\delta r\\ &= \frac{-r_s}{K}\left[s\delta s + s\alpha_{rs}\delta r + (\delta s)^2 + \alpha_{rs}\delta s\delta r\right] - t_s\delta s + t_r\delta r\\ &\simeq \frac{-r_s}{K}\left[s(\delta s+\alpha_{rs}\delta r)\right] - t_s\delta s + t_r\delta r \text{ (Neglecting second order terms)}\end{aligned}$$

Setting $s = s^*$ from Equation S2,

$$\begin{aligned}\dot{s}|_{(s+\delta s,\ r+\delta r)} &= \frac{-r_s(\alpha_{rs}-1)}{(\alpha_{rs}\alpha_{sr}-1)}(\delta s+\alpha_{rs}\delta r) - t_s\delta s + t_r\delta r\\ &= \delta s\left[\frac{r_s(1-\alpha_{rs})}{\alpha_{rs}\alpha_{sr}-1} - t_s\right] + \delta r\left[\frac{r_s\alpha_{rs}(1-\alpha_{rs})}{\alpha_{rs}\alpha_{sr}-1} + t_r\right]\end{aligned} \tag{S6}$$

Due to symmetry, $\dot{r}$ for a small perturbation $(\delta s,\ \delta r)$ about the fixed point $(s,\ r)$ is given by

$$\dot{r}|_{(s+\delta s,\ r+\delta r)} = \delta r\left[\frac{r_r(1-\alpha_{sr})}{\alpha_{rs}\alpha_{sr}-1} - t_r\right] + \delta s\left[\frac{r_r\alpha_{sr}(1-\alpha_{sr})}{\alpha_{rs}\alpha_{sr}-1} + t_s\right] \tag{S7}$$

The stability of the fixed point $(s,\ r)$, as given by Equation S2, can be inferred based on the signs of $\dot{s}$ and $\dot{r}$ in Equations S6 and S7 for different combinations of $\delta s$, $\delta r$, as well as the magnitude of $\alpha_{rs}$ and $\alpha_{sr}$. Two sets of perturbations are possible, which can be visualised in an $s$-$r$ plane with the fixed point as the origin. When $\delta s$ and $\delta r$ are either both positive or both negative, these perturbations correspond to the first and third quadrant respectively, while when one is positive and the other negative, the perturbations lie in the second and fourth quadrants. We will now use Equations S6 and S7 to consider the behaviour of $\dot{s}$ and $\dot{r}$ in each of these cases to infer the stability of the fixed point.

**For $\delta s,\ \delta r > 0$:**
When $\alpha_{rs},\ \alpha_{sr} > 1$, $1-\alpha_{rs}\alpha_{sr} < 0$, so the coefficient of $\delta s$ in Equation S6 is always negative. For the fixed point to be stable, we require that $\dot{s}$ and $\dot{r}$ are both negative for positive $\delta s$ and $\delta r$. From Equation S5, we see that $t_r < \left|\dfrac{r_s\alpha_{rs}(1-\alpha_{rs})}{\alpha_{rs}\alpha_{sr}-1}\right|$ implies $\dot{s} < 0$, and by symmetry, $t_s < \left|\dfrac{r_r\alpha_{sr}(1-\alpha_{sr})}{\alpha_{rs}\alpha_{sr}-1}\right|$ implies $\dot{r} < 0$. For simplicity, we denote $E_s = \dfrac{r_s\alpha_{rs}(1-\alpha_{rs})}{\alpha_{rs}\alpha_{sr}-1}$ and $E_r = \dfrac{r_r\alpha_{sr}(\alpha_{sr}-1)}{1-\alpha_{rs}\alpha_{sr}}$.

These two inequalities are also sufficient conditions for the stability of the fixed point when $\alpha_{rs},\ \alpha_{sr} < 1$. Here, $\alpha_{rs}-1, \alpha_{sr-1} < 0$ while $1-\alpha_{rs}\alpha_{sr} > 0$ so that the respective first term in Equations S6 and S7 is still negative.

For $\alpha_{rs} < 1,\ \alpha_{sr} > 1$, and $\alpha_{rs}\alpha_{sr} > 1$, the $\delta r$ term in Equation S6 is always positive. Therefore, $\dot{s}$ is negative only if:

1. $t_s > \left|\dfrac{E_s}{\alpha_{rs}}\right|$, and

2. $\left|\delta s\left[\frac{E_s}{\alpha_{rs}} - t_s\right]\right| > |\delta r(E_s + t_r)|$

The first inequality above is never satisfied in our parameterisation of the model since $t_s$ is at least an order of magnitude less than $r_s$ and all the $\alpha$ values lie between 0 and 7.4. Furthermore, for the same conditions on the $\alpha$ values, the $\delta r$ term in Equation S7 is always negative. For $\dot{r} < 0$ then, we require $t_s < |E_r|$. The two inequalities in $t_s$ are contradictory, and the co-existence fixed point is therefore unstable for $\alpha_{rs} < 1,\ \alpha_{sr} > 1$, and $\alpha_{rs}\alpha_{sr} > 1$. A similar set of contradictory inequalities in $t_r$ can be derived for $\alpha_{rs} < 1,\ \alpha_{sr} > 1$, and $\alpha_{rs}\alpha_{sr} < 1$, indicating that the fixed point is unstable. This can further be extended to show that the fixed point is also unstable for $\alpha_{rs} > 1,\ \alpha_{sr} < 1$. The fact that a feasible condition exists in each of the above cases for only one species to satisfy the requirements of stability could also indicate behaviour similar to the steady state behaviour of the two-species LV model without transitions, resulting in single-species steady states with a clear winner (System S1).

**For $\delta s,\ \delta r < 0$:**

In this case, we require $\dot{s},\ \dot{r} > 0$ for the fixed point to be stable. When $\alpha_{rs},\ \alpha_{sr} > 1$, $(1 - \alpha_{rs}\alpha_{sr}) < 0$ and the $\delta s$ term in Equation S6 is always positive. Therefore, $\dot{s} > 0$ if $t_r < |E_s|$, and likewise, $\dot{r} > 0$ if $t_s < |E_r|$. As earlier, the same inequalities hold for $\alpha_{rs},\ \alpha_{sr} < 1$.

For $\alpha_{rs} < 1,\ \alpha_{sr} > 1$, while $\alpha_{rs}\alpha_{sr} < 1$, the $\delta s$ term in Equation S6 is always positive. $\dot{s}$ is therefore positive if $t_r < |E_s|$, and the inequality from earlier continues to hold for negative perturbations from the fixed point. But as derived earlier for positive perturbations, it can be shown that for $\alpha_{rs}\alpha_{sr} > 1$, the fixed point cannot be stable in our parameterisation of the model.

Taken together, for positive and negative perturbations of both $s$ and $r$ i.e., the first and third quadrants of the $s$-$r$ plane, the fixed point is stable if:

1. $t_r < |E_s|$ and $t_s < |E_r|$, and
2. $\alpha_{rs},\ \alpha_{sr} > 1$, or $\alpha_{rs},\ \alpha_{sr} < 1$

**For $\delta s < 0,\ \delta r > 0$ or $\delta s > 0,\ \delta r < 0$:**

These perturbations lie in the second and fourth quadrants of the $s$-$r$ plane respectively. For stability in the second quadrant, where $\delta s < 0$ and $\delta r > 0$, we require $\dot{s} > 0$ and $\dot{r} < 0$. For $\alpha_{rs},\ \alpha_{sr} > 1$, the $\delta s$ term in Equation S6 is always positive. Therefore, for $\delta r > 0$, $\dot{s}$ will be positive if $t_r > |E_s|$. Likewise, we require $\dot{r}$ to be negative for the fixed point to be stable, for which, from Equation S7, we require $t_s > |E_r|$. As explained earlier, both these inequalities are never satisfied in our parameterisation of the model, which indicates that the fixed point is unstable for $\delta s < 0,\ \delta r > 0$. As in the previous case, this can be extended to $\alpha_{rs},\ \alpha_{sr} < 1$ to show that the fixed point is unstable in the second and fourth quadrants of the $s$-$r$ plane.

For $\alpha_{rs} < 1,\ \alpha_{sr} > 1$ and $\alpha_{rs}\alpha_{sr} > 1$, the $\delta r$ term in Equation S6 is always positive. Therefore, for $\dot{s} > 0$, we require $t_s > \left|\frac{E_s}{\alpha_{rs}}\right|$. Likewise, for the same conditions on the $\alpha$s, the $\delta r$ term in Equation S7 is always negative. Therefore, for $\dot{r} < 0$, we require $t_s > |E_r|$. Neither of these inequalities is satisfied in our parameterisation of the extended System S3.

For $\alpha_{rs} < 1,\ \alpha_{sr} > 1$ and $\alpha_{rs}\alpha_{sr} < 1$, the $\delta s$ term in Equation S6 is always positive. Therefore, for $\dot{s} > 0$, we require $t_r > |E_s|$. For the same conditions, the $\delta s$ in Equation S7 is always negative. Therefore, for $\dot{r} < 0$, we require $t_r > \left|\frac{E_r}{\alpha_{sr}}\right|$. As explained earlier, neither of these inequalities is satisfied in our model, which indicates that the fixed point is unstable for $\alpha_{rs} < 1,\ \alpha_{sr} > 1$ and $\alpha_{rs}\alpha_{sr} < 1$. Similar inequalities can also be derived for $\alpha_{rs} > 1,\ \alpha_{sr} < 1$, both in the second quadrant as shown here, and in the fourth quadrant where $\delta s > 0,\ \delta r < 0$. Taken together, this indicates that the fixed point is unstable for all possible configurations of $\alpha_{rs}$ and $\alpha_{sr}$ in the second and fourth quadrants of the $s$-$r$ plane.

Considering the whole $s$-$r$ plane over all the parameter combinations, we see that the fixed point from the two-species LV model is stable in the first and third quadrants, and unstable in the other two, for $\alpha_{rs},\ \alpha_{sr} > 1$, or $\alpha_{rs},\ \alpha_{sr} < 1$, and unstable in all four quadrants for any other parameter combinations. This indicates that it is a saddle point for $\alpha_{rs},\ \alpha_{sr} > 1$, or $\alpha_{rs},\ \alpha_{sr} < 1$, with the separatrix given by the inequalities in $t_r$ and $t_s$ derived above.

#### S3 Alternate view of steady state behaviour

The foregoing analyses build a picture of the steady state behaviour of the extended System S3, based on what is known about the simpler two-species LV model. In this final section, we motivate an alternative analytical view that could potentially provide an additional numerical approach regarding the full steady state behaviour of System S3 without relying on the known properties of the LV system.

To begin with, let us consider once again, the extended System S3. The steady state of the total population can be expressed as $\dot{s} + \dot{r} = 0$. The case of both individual populations individually being at steady state is a specific case of the total population equilibrium where $\dot{s} = 0$ and $\dot{r} = 0$.

**For $\dot{s} + \dot{r} = 0$:**

System S3 is a closed system. Specifically, the transition terms cancel out when adding the two equations, yielding

$$\dot{s} + \dot{r} = r_s s - \frac{r_s s^2}{K} - \frac{r_s \alpha_{rs} sr}{K} + r_r r - \frac{r_r r^2}{K} - \frac{r_r \alpha_{sr} sr}{K} = 0$$

$$\implies -\frac{r_s}{K} s^2 - \frac{r_s \alpha_{rs} + r_r \alpha_{sr}}{K} sr - \frac{r_r}{K} r^2 + r_s s + r_r r = 0 \tag{S8}$$

Equation S8 represents a family of conic sections in the $s$-$r$ plane of the form $As^2 + Bsr + Cr^2 + Ds + Er = 0$, where $A = -\frac{r_s}{K}$, $B = -\frac{r_s \alpha_{rs} + r_r \alpha_{sr}}{K}$, $C = -\frac{r_r}{K}$, $D = r_s$, and $E = r_r$. The discriminant of this equation, given by $\nabla = B^2 - 4AC$ will determine its shape, such that Equation S8 is an ellipse for $\nabla < 0$, a parabola for $\nabla = 0$, and a hyperbola for $\nabla > 0$. For this system,

$$\nabla = \frac{(r_s \alpha_{rs} + r_r \alpha_{sr})^2}{K^2} - \frac{4 r_s r_r}{K^2}$$

Therefore, Equation S8 forms an ellipse if $r_s \alpha_{rs} + r_r \alpha_{sr} > 2\sqrt{r_s r_r}$, a parabola if $r_s \alpha_{rs} + r_r \alpha_{sr} = 2\sqrt{r_s r_r}$, and a hyperbola if $r_s \alpha_{rs} + r_r \alpha_{sr} < 2\sqrt{r_s r_r}$. For the special case where $r_s = r_r = r$, these conditions can be simplified further such that the final shape depends only on the values of $\alpha_{rs}$ and $\alpha_{sr}$.

Equation S7 represents a family of curves for which the sum $\dot{s} + \dot{r}$ is zero. The intersection of these curves with either $\dot{s} = 0$ or $\dot{r} = 0$ can therefore be evaluated numerically to compute trajectories in the $s$-$r$ plane that correspond to the full steady state behaviour of System S3. Such comprehensive analysis is beyond the scope of our current work, but provides a method of systematic comparison for identifying general co-existence equilibria when they do not overlap with classical results from the LV model.
