## Supplementary Tables for "Impacts of competition and phenotypic plasticity on the viability of adaptive therapy"

### Supplementary Table 1: Pairwise Wilcoxon’s signed-rank tests comparing steady state population sizes across model types

```
popsize_df <- no_therapy %>%
    select(ModelType, PopSizeSS)

pairwise.wilcox.test(popsize_df$PopSizeSS, popsize_df$ModelType, p.adjust.method = "BH",
    paired = F)
```

```
##
##  Pairwise comparisons using Wilcoxon rank sum test with continuity correction
##
## data:  popsize_df$PopSizeSS and popsize_df$ModelType
##
##       AC     AC+Tr  SC
## AC+Tr <2e-16 -      -
## SC    <2e-16 <2e-16 -
## SC+Tr <2e-16 <2e-16 <2e-16
##
## P value adjustment method: BH
```

### Supplementary Table 2: Mean differences in number of “Favourable” outcomes between dose levels under constant dose therapy (CDT)

```
## # A tibble: 4 x 4
## # Groups:   ModelType [4]
##   ModelType contrast                  MeanDiff Error
##   <fct>     <chr>                        <dbl> <dbl>
## 1 AC        Cytostatic2 - Cytostatic1    1467.  19.9
## 2 AC+Tr     Cytostatic2 - Cytostatic1     889.  45.9
## 3 SC        Cytostatic2 - Cytostatic1     938   30.8
## 4 SC+Tr     Cytostatic2 - Cytostatic1     362.  56.0
```

[*]Equal contribution
[†]Equal contribution
[‡]Corresponding author
[§]Corresponding author

### Supplementary Table 3: Pairwise Wilcoxon's signed-rank tests comparing cycling time periods across model types

```
cycling_df <- at_df %>%
    filter(Outcome == "Cycling") %>%
    select(ModelType, Period)

pairwise.wilcox.test(cycling_df$Period, cycling_df$ModelType, p.adjust.method = "BH",
    paired = F)
```

```
##
##  Pairwise comparisons using Wilcoxon rank sum test with continuity correction
##
## data:  cycling_df$Period and cycling_df$ModelType
##
##       AC      AC+Tr   SC
## AC+Tr 2.2e-14 -       -
## SC    0.5574  1.3e-12 -
## SC+Tr < 2e-16 0.0059  < 2e-16
##
## P value adjustment method: BH
```

### Supplementary Table 4: PC loadings and component variances for resistant cell fraction before and after constant dose therapy

```
pca.cdt <- prcomp(~r_x + r_y + c + d + a + b + ResFracBT + ResFracCDT, data = cdt_pca_df,
    scale. = T)
pca.cdt
```

```
## Standard deviations (1, .., p=8):
## [1] 1.4687929 1.2275332 1.1212790 1.0893080 0.9010804 0.7165986 0.6466216
## [8] 0.3851911
##
## Rotation (n x k) = (8 x 8):
##                    PC1         PC2         PC3         PC4         PC5
## r_x         0.23519914  0.04136956 -0.46650405  0.48281677  0.60676942
## r_y         0.27848800  0.03211851  0.53597643 -0.22923302  0.68121893
## c           0.06458232 -0.70900496 -0.12355626 -0.09558939  0.04588543
## d           0.01787494 -0.69674464  0.15908973  0.19024430 -0.02180195
## a           0.46679445  0.06861358 -0.06063585  0.53061097 -0.28820452
## b           0.59129314  0.04843832  0.15785644 -0.18464893 -0.23629943
## ResFracBT  -0.10573659  0.03062638  0.65008624  0.54378746 -0.09341779
## ResFracCDT -0.53285134  0.03352268  0.05761995  0.24012897  0.13253190
##                   PC6         PC7        PC8
## r_x         0.1683283 -0.02507731 -0.3083820
## r_y        -0.1909456 -0.01480881  0.2843512
## c          -0.4027398  0.52137617 -0.1805537
## d           0.3355784 -0.55948682  0.1631297
## a          -0.3912807  0.01663941  0.5056986
## b          -0.1942276 -0.34543692 -0.6132893
## ResFracBT   0.1886111  0.35226685 -0.3180753
## ResFracCDT -0.6587823 -0.41303479 -0.1783456
```

```r
summary(pca.cdt)
```

```
## Importance of components:
##                           PC1    PC2    PC3    PC4    PC5     PC6     PC7
## Standard deviation     1.4688 1.2275 1.1213 1.0893 0.9011 0.71660 0.64662
## Proportion of Variance 0.2697 0.1883 0.1572 0.1483 0.1015 0.06419 0.05226
## Cumulative Proportion  0.2697 0.4580 0.6152 0.7635 0.8650 0.92919 0.98145
##                            PC8
## Standard deviation     0.38519
## Proportion of Variance 0.01855
## Cumulative Proportion  1.00000
```

#### Supplementary Table 5: PC loadings and component variances for total population size before and after constant dose therapy

```r
pca.cdt <- prcomp(~r_x + r_y + c + d + a + b + PopSizeBT + PopSizeCDT, data = cdt_pca_df,
    scale. = T)
pca.cdt
```

```
## Standard deviations (1, .., p=8):
## [1] 1.4585649 1.3369177 1.1416813 1.0007530 0.8476204 0.6840118 0.6163875
## [8] 0.4626348
##
## Rotation (n x k) = (8 x 8):
##                    PC1         PC2        PC3        PC4          PC5
## r_x         0.13221204 -0.37259380  0.4253302 -0.6214518  0.006658186
## r_y         0.27327142 -0.18145839 -0.6732715 -0.2339408 -0.252192899
## c          -0.50402714 -0.15842466 -0.1970170 -0.2981196 -0.238284136
## d          -0.47133477 -0.12003634 -0.2629306 -0.3779468  0.238371354
## a           0.07710987 -0.61097394  0.1627244  0.1141547  0.503880363
## b           0.09985118 -0.56778644 -0.3209707  0.4227107 -0.062466946
## PopSizeBT   0.54184437 -0.06938035  0.0843523 -0.2669794 -0.370666211
## PopSizeCDT  0.34947299  0.29707808 -0.3481880 -0.2536861  0.653883169
##                    PC6         PC7         PC8
## r_x         0.01329567 -0.44486996 -0.28009886
## r_y         0.05951467 -0.35896998  0.43401619
## c           0.63171588  0.33108961 -0.16644128
## d          -0.67336555  0.20052959  0.03169160
## a           0.20080163  0.24395753  0.47699307
## b          -0.13636825 -0.01775528 -0.60260108
## PopSizeBT  -0.18170128  0.67267274 -0.01733063
## PopSizeCDT  0.22773265  0.10526004 -0.33692243
```

```r
summary(pca.cdt)
```

```
## Importance of components:
##                           PC1    PC2    PC3    PC4     PC5     PC6     PC7
## Standard deviation     1.4586 1.3369 1.1417 1.0008 0.84762 0.68401 0.61639
## Proportion of Variance 0.2659 0.2234 0.1629 0.1252 0.08981 0.05848 0.04749
## Cumulative Proportion  0.2659 0.4894 0.6523 0.7775 0.86727 0.92575 0.97325
##                            PC8
## Standard deviation     0.46263
## Proportion of Variance 0.02675
## Cumulative Proportion  1.00000
```

#### Supplementary Table 6: PC loadings and component variances for resistant cell fraction before and after adaptive therapy

```
pca.at <- prcomp(~r_x + r_y + c + d + a + b + ResFracBT + ResFracAT, data = at_pca_df,
    scale. = T)
pca.at
```

```
## Standard deviations (1, .., p=8):
## [1] 1.3924538 1.2497326 1.1468431 1.0551435 0.9069109 0.8228966 0.6402570
## [8] 0.4013589
##
## Rotation (n x k) = (8 x 8):
##                   PC1         PC2         PC3         PC4         PC5
## r_x         0.16328243 -0.24187517  0.58188727 -0.27785936 -0.50015482
## r_y         0.34593644 -0.07144851 -0.41228796  0.25849090 -0.73267238
## c           0.16414210  0.58899661  0.33474395  0.16221460 -0.12140106
## d           0.23539114  0.63777793  0.09622747 -0.21916777 -0.04656261
## a           0.47401081 -0.30368494  0.28625691 -0.35462112  0.23775354
## b           0.56758135 -0.22906056 -0.04337974  0.32950320  0.21578807
## ResFracBT   0.08008605  0.06107036 -0.48851353 -0.73841314 -0.08100090
## ResFracAT  -0.46697602 -0.18517816  0.21827121 -0.04523359 -0.29407085
##                   PC6         PC7         PC8
## r_x         0.39277737  0.02737459  0.30636311
## r_y        -0.08698881 -0.03113967 -0.30508791
## c          -0.34241102 -0.56852723  0.18046204
## d           0.01547869  0.67283210 -0.15945341
## a          -0.32266402 -0.13933866 -0.54344824
## b          -0.26378934  0.28646693  0.56277518
## ResFracBT  -0.17682728 -0.16964116  0.37329618
## ResFracAT  -0.71812898  0.30347655  0.06057347
```

```
summary(pca.at)
```

```
## Importance of components:
##                           PC1    PC2    PC3    PC4    PC5     PC6     PC7
## Standard deviation     1.3925 1.2497 1.1468 1.0551 0.9069 0.82290 0.64026
## Proportion of Variance 0.2424 0.1952 0.1644 0.1392 0.1028 0.08464 0.05124
## Cumulative Proportion  0.2424 0.4376 0.6020 0.7412 0.8440 0.92862 0.97986
##                            PC8
## Standard deviation     0.40136
## Proportion of Variance 0.02014
## Cumulative Proportion  1.00000
```

#### Supplementary Table 7: PC loadings and component variances for total population size before and after adaptive therapy

```
pca.at <- prcomp(~r_x + r_y + c + d + a + b + PopSizeBT + PopSizeAT, data = at_pca_df,
    scale. = T)
pca.at
```

```
## Standard deviations (1, .., p=8):
## [1] 1.4783901 1.4180084 1.0993056 1.0349681 0.7324225 0.6567598 0.5363972
## [8] 0.5181552
##
```

```
## Rotation (n x k) = (8 x 8):
##                   PC1         PC2         PC3         PC4         PC5
## r_x        0.1020732  0.25560768 -0.71484015  0.36098269 -0.01819722
## r_y        0.1685799  0.24484616  0.49515177  0.63740091  0.06375176
## c          0.1952774 -0.47516204 -0.10992927  0.40849129  0.52565813
## d          0.2406283 -0.48126260 -0.09182357  0.33598674 -0.65245833
## a          0.4979006  0.21748538 -0.36404174 -0.18645836  0.19721522
## b          0.5413589  0.21397863  0.28069853 -0.05059909  0.20000223
## PopSizeBT -0.1008167  0.56757004 -0.01918980  0.25698421 -0.31116713
## PopSizeAT -0.5603465  0.04582496 -0.10782381  0.28335751  0.34331693
##                    PC6          PC7         PC8
## r_x        0.22585485  0.477582561 -0.05928057
## r_y        0.30087734 -0.096643346 -0.39527783
## c         -0.52975203  0.008517187 -0.01058137
## d          0.14137530 -0.240660345  0.29250547
## a          0.10745182 -0.686002128 -0.12830964
## b          0.09971616  0.272414126  0.67500972
## PopSizeBT -0.65959817 -0.181401113  0.19102362
## PopSizeAT  0.31824403 -0.356188541  0.49596202
```

```
summary(pca.at)
```

```
## Importance of components:
##                           PC1    PC2    PC3    PC4     PC5     PC6     PC7
## Standard deviation     1.4784 1.4180 1.0993 1.0350 0.73242 0.65676 0.53640
## Proportion of Variance 0.2732 0.2513 0.1511 0.1339 0.06706 0.05392 0.03597
## Cumulative Proportion  0.2732 0.5245 0.6756 0.8095 0.87656 0.93047 0.96644
##                            PC8
## Standard deviation     0.51816
## Proportion of Variance 0.03356
## Cumulative Proportion  1.00000
```
