## Supplementary Figures for "Impacts of competition and phenotypic plasticity on the viability of adaptive therapy"

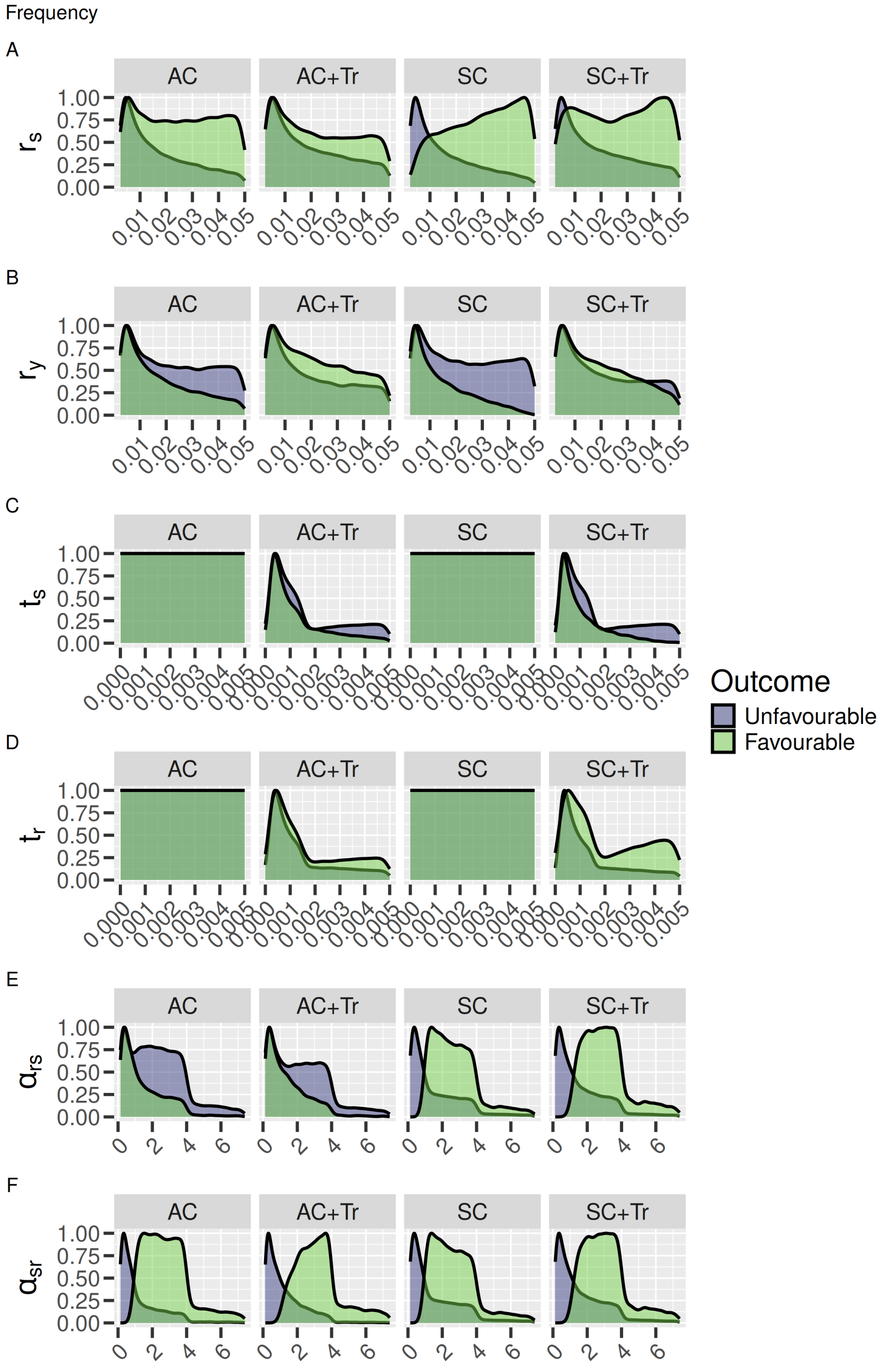


Supplementary Figure S1: Distribution of all parameter values without any therapy across model types. Favourable indicates $s \geq 0.85$ at steady. Parameter notations and model type definitions are given in the main text.

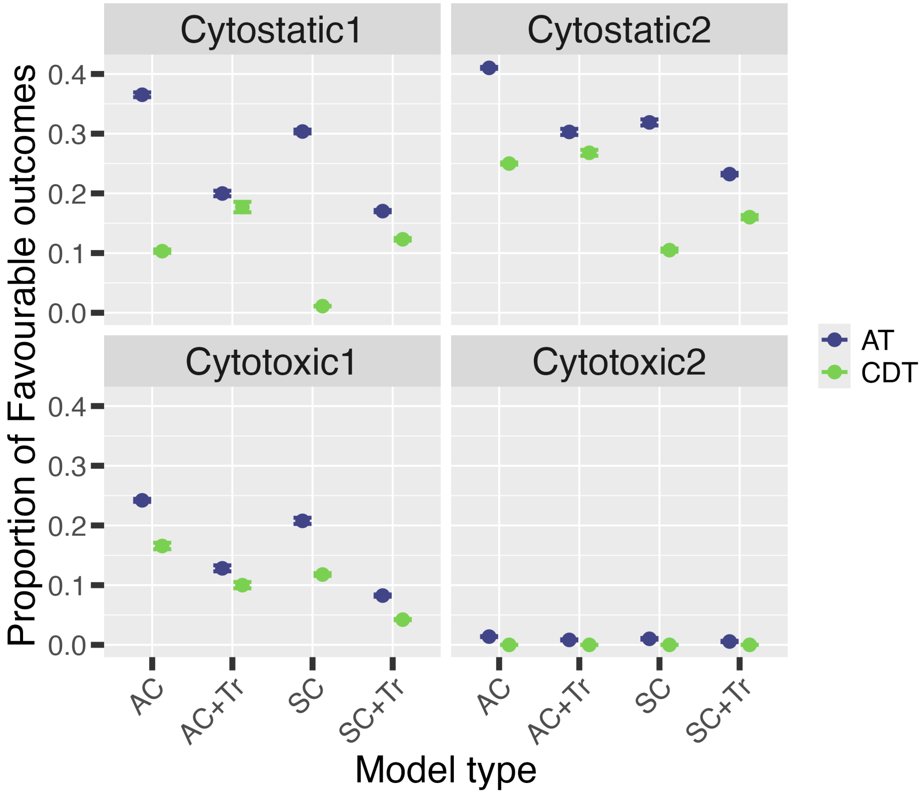


Supplementary Figure S2: Fraction of "Favourable" outcomes across model types for constant dose therapy (CDT) vs adaptive therapy (AT). Cytostatic1 and Cytostatic2 correspond to $\gamma = 0.33$ and 0.1 in System 2, ad Cytotoxic1 and Cytotoxic2 correspond to $d_{th} = 0.01$ and 0.05 resprectively. Points and error bars are mean $\pm$ SD over all replicate runs.
